# Conformational plasticity of LptC regulates lipopolysaccharide transport by the LptB_2_FGC complex

**DOI:** 10.1101/2025.07.09.663713

**Authors:** Aaron Klausnitzer, Jagdeep Kaur, Tobias Rath, Samuel Seidl, Johanna Becker-Baldus, Nina Morgner, Clemens Glaubitz

## Abstract

The outer membrane of Gram-negative bacteria is coated with lipopolysaccharide (LPS). The Lpt system maintains membrane asymmetry by transporting LPS from the inner to the outer membrane. Transport begins with the LptB₂FGC complex, where the ABC transporter LptB₂FG associates with LptC to extract LPS. LPS is then passed via LptA to the LptDE translocon. While LptB₂FGC structures suggest an extrusion mechanism, the role of LptC remains unclear. Here, we reconstituted the complex in vitro from purified LptB₂FG and LptC, and demonstrate that LptC stabilizes the complex and modulates ATPase activity. Using differential isotope labeling and solid-state NMR including dynamic nuclear polarization, we observed that the LptC transmembrane helix LptC_TMH_ is tightly associated with the transporter in the apo state. Upon LPS or ATP binding, LptC_TMH_ becomes dynamic, favoring cavity collapse and substrate-coupled ATPase activity. Our data support a model in which LptC acts as a mechanical transducer linking transport and energy consumption.

## Introduction

Gram-negative bacteria have developed a unique way of encapsulation, which features a symmetric inner membrane (IM) and an asymmetric outer membrane (OM). The IM and the inner leaflet of the OM contain phospholipids, whereas the outer leaflet of the OM is almost exclusively composed of lipopolysaccharides (LPS) (*1–3*). Outer membrane biogenesis is driven by modification, transport and embedding LPS in the OM, which plays an important role in developing resistance mechanisms against antibiotics (*2, 4, 5*). Therefore, the lipopolysaccharide transport (Lpt) machinery has shifted into the focus of developing potent antibiotic compounds, which are urgently needed to overcome bacterial multiresistance (*6–11*). This cell envelope spanning transport bridge, which is assembled by the seven proteins LptA, B, C, D, E, F and G, pushes LPS molecules from the inner to the outer membrane in a PEZ-dispenser-like mechanism (Fig. 1A) (*12, 13*). The bridge is anchored in the IM by LptB_2_FG, a type VI ATP-binding cassette (ABC) transporter, which extracts LPS from the IM by ATP binding and hydrolysis (*14–16*). The Lpt bridge itself is formed by at least one copy of LptA. They are connected via β-jellyroll-like (βJR) domains to LptF, LptC at the IM and to the outer membrane translocon LptDE, which inserts the transferred LPS into the OM (*17–20*).

**Figure 1:**
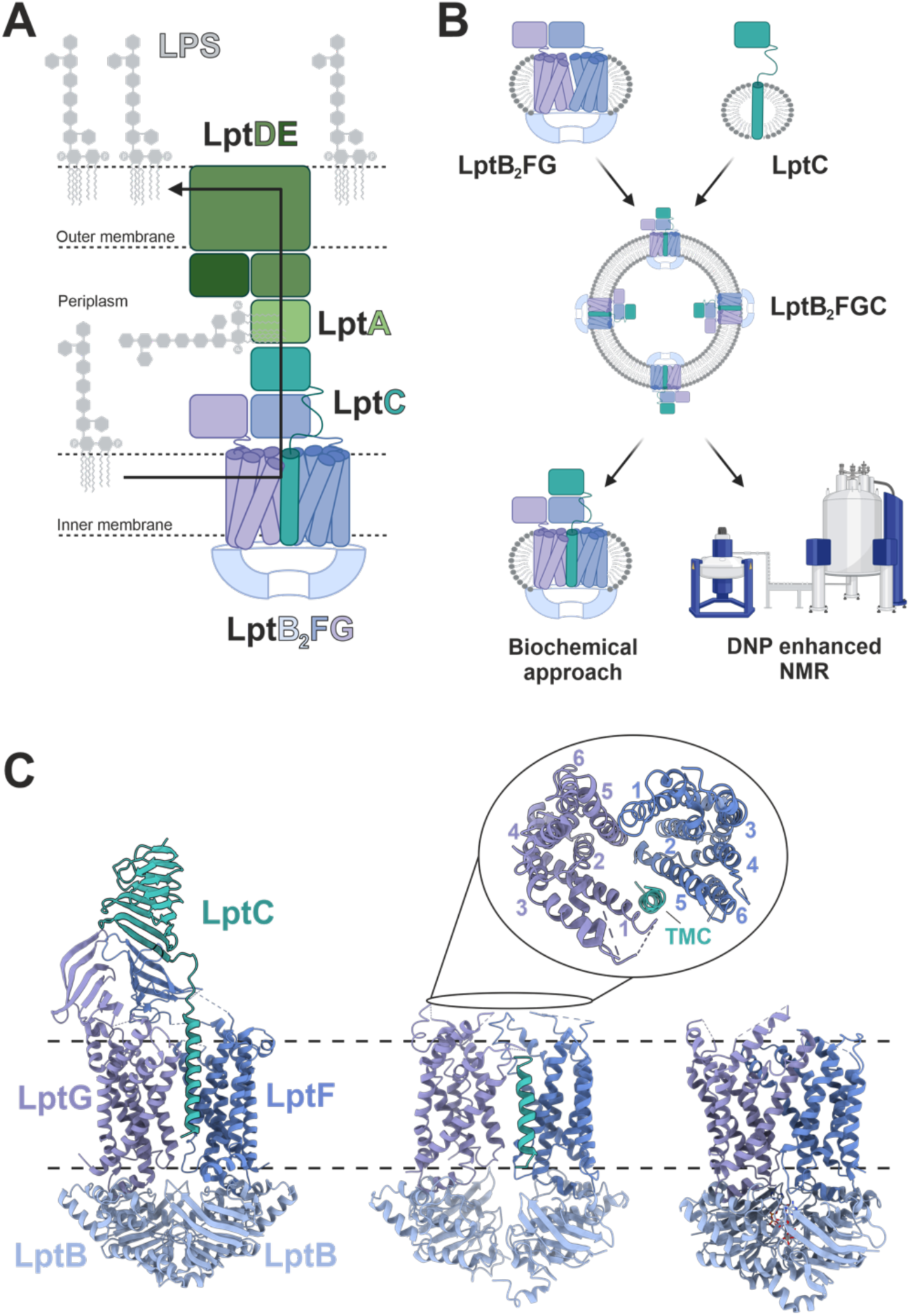
Lipopolysaccharide transport across the cell envelope. **(A)** Schematic overview of the envelope spanning lipopolysaccharide transport system (PDBs: 6MJP, 2R1A and 7OMM). LptB2FGC anchors the bridge at the IM, while LptDE provides the contact-site with the OM. Both are connected via the multiple copies of soluble LptA. LPS transport across the protein bridge is indicated by the black arrow. The peptidoglycan layer was omitted for simplification. **(B)** Overview of the experimental pipeline for the investigation of the inner membrane LPS transporter. LptB2FG and LptC are expressed separately for differential NMR labelling and co-reconstituted into liposomes. Experimental data was recorded with the re-solubilized LptB2FGC as well as LptB2FGC in liposomes. **(C)** Crystal structure of LptB2FGC in the nucleotide-free state (PDB: 6MJP), Cryo-EM structures of the nucleotide free (PDB:6MI7) and vanadate-trapped (ADP-VO4-trapped) LptB2FGC complex (PDB: 6MHZ). Figure (A) and (B) were created using BioRender (*51*).

The ABC transporter LptB_2_FGC contains the homodimeric nucleotide binding domains (NBDs) LptB, the heterodimeric transmembrane domains (TMDs) LptF and LptG as well as LptC, which is a βJR anchored within the TMDs via a hydrophobic α-helix of debated functionality. Crystal structures of the LptB_2_FGC complex in the LPS bound state show the two transmembrane domains (TMDs) LptF and LptG in a pseudo-two-fold symmetry (Fig. 1C). Both enclose a V-shaped hydrophobic cavity that forms the LPS binding pocket. The single transmembrane helix of LptC (LptC_TMH_) associates with the TMDs of LptFG and is sandwiched between the TMH 1 of LptG and TMH 5 of LptF. The βJR of LptC associates with the βJR of LptF building a continuous groove surrounding the acyl chains of LPS after translocation. (*21*) Recent structural Cryo-EM snapshots of LptB_2_FGC in the ATP-bound (AMP-PNP-bound) and high energy transition states (ADP-VO_4_-trapped) suggest an extrusion mechanism in which the V-shaped cavity of LptFG collapses and pushes LPS out of the TMDs (*16*). However, the AMP-PNP-bound and ADP-VO₄-trapped structures lack sufficient resolution to resolve the jellyroll domains of LptF, LptG, LptC, and the transmembrane helix (TMH) of LptC. Based on these cryo-EM structures of LptB₂FGC, it is proposed that the TMH of LptC is displaced from the transporter upon collapse of the transmembrane domains (TMDs) (*15, 16*).

Uncovering the role of LptC in LPS transport has become a topic of interest as its presence appears to be a unique feature of this ABC transporter complex (*22*). Although essential for LPS transport, biochemical studies have shown that the association of LptC with the LptB₂FG complex reduces its ATPase activity by approximately 50% (*15, 16, 21*). This effect has been shown to be dependent on LptC_TMH_ association with LptB_2_FG (*23*). Interestingly, in the absence of the LptC TMH, LptB₂FG can still associate with the soluble β-jellyroll (βJR) domain of LptC and mediate LPS transport (*17*). Furthermore, a single point mutation in the βJR of LptF has been shown to bypass the requirement for LptC altogether (*24, 25*). These findings raise an important question: if only the βJR of LptC is necessary for function, why is the full structure of LptC—including its TMH—conserved among orthologous proteins (*26*).

In exploring its potential role, LptC_TMH_ has been shown to be important for anchoring the protein in the IM (*27*). A loss-of-function (LOF) mutant screen further underscored the significance of LptC_TMH_, revealing that multiple residues within the helix are critical for maintaining outer membrane (OM) stability. Additionally, some of these LptC_TMH_ mutants in the LptB_2_FGC complex exhibit altered ATPase activities (*23*). Despite these insights, the precise function of LptC_TMH_ in LPS transport - as well as the mechanisms underlying its displacement and dynamics within the ABC transporter—remains unclear.

To investigate the role of LptC, we developed a protocol for in vitro assembly of the active LptB₂FGC transporter complex from purified LptB₂FG and LptC. Our results show that LptC stabilizes the complex and modulates both its basal and LPS-stimulated ATPase activity. This in vitro reconstitution enabled the application of differential isotope labeling strategies for both ambient-temperature and cryogenic DNP-enhanced solid-state NMR (ssNMR) spectroscopy on the LptB₂FGC complex reconstituted in liposomes (Fig. 1B). Using this approach, we demonstrate that LptC_TMH_ is tightly associated with LptB_2_FG in the apo state. Upon LPS binding and/or formation of the ATPase high-energy transition state, the system exhibits increased dynamics with LptC_TMH_ exchanging between two distinct conformations. These findings are further supported by native mass spectrometry and support a model in which LptC functions as a mechanical transducer, coupling LPS transport to the ATPase activity of the Lpt machinery.

## Results

### *In vitro* complex formation and LPS stimulation of LptB_2_FGC

The Lpt system exhibits inherent modularity (*28, 29*), which allowed us to express LptB_2_FG and LptC separately for targeted NMR labeling strategies. While the association of LptB_2_FG and LptC has been demonstrated *in vivo* and *in situ*, there are no reports describing the assembly of the complex *in vitro* (*17, 30*). Notably, formation of the LptB_2_FGC complex has been associated with reduced ATPase activity, as shown in *in vitro* assays with the *in vivo*-assembled complex (*13, 15, 16*). To directly examine complex assembly in vitro, we developed a co-reconstitution protocol incorporating both full-length proteins into liposomes. We separately expressed and purified LptB_2_FG and LptC in detergent micelles and assembled the complex in a native lipid environment by co-reconstituting both into POPE/POPG liposomes. Proper complex formation was confirmed by monitoring the reduction in ATPase activity of LptB₂FG as a function of the LptC:LptB₂FG molar ratio (Fig. 2A). Increasing amounts of LptC led to a progressive decrease in ATPase activity, reaching a minimum of approximately 45% at a two-fold molar excess of LptC. To further explore this regulatory function of LptC we introduced LPS into proteoliposomes containing either LptB_2_FG or LptB_2_FGC. For the LptB₂FGC complex, a LptC:LptB₂FG molar ratio of 2:1 was used to ensure complex saturation. We then compared the stimulation of the ATPase activity in response to LPS titration (Fig 2B). Substrate binding has been shown to stimulate the ATPase activity of various ABC transporters (*31–37*). Both LptB_2_FG and LptB_2_FGC exhibited a 2 - 2.5 fold increase in ATPase activity upon LPS addition. However, the absolute ATPase activity of LptB₂FGC remained significantly lower than that of LptB₂FG alone (see Materials and Methods). This inhibitory effect is consistent with previous reports and confirms that we successfully reconstituted a functional LptB₂FGC complex *in vitro* (*15, 16*).

**Figure 2:**
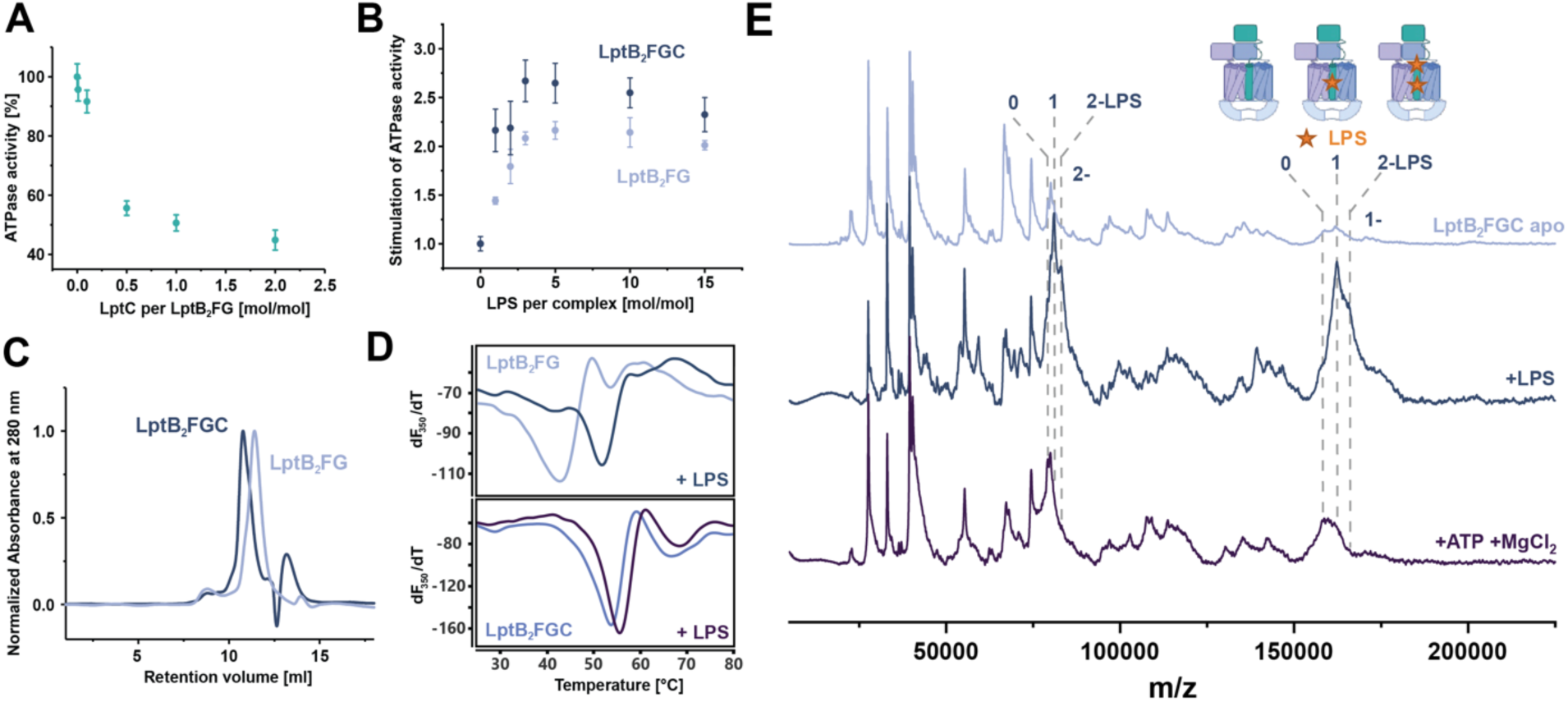
*In vitro* complex formation and re-solubilisation of LptB2FGC. **(A)** ATPase activity under titration of LptC to LptB2FG in liposomes. Activities are shown in percent relative to the activity of LptB2FG alone in liposomes. Each data point represents the averaged Vmax value ± standard error as determined from triplicate ATPase assays. **(B)** ATPase activity under titration of LPS to LptB2FG and LptB2FGC in liposomes. Activities are shown relative to the activity of LptB2FG and LptB2FGC in liposomes in the absence of LPS. Each data point represents the averaged Vmax value ± standard error as determined from triplicate ATPase assays. **(C)** Size exclusion chromatograms of LptB2FG and assembled LptB2FGC (resolubilized from liposomes) in DDM micelles normalized to the respective peak maximum. **(D)** NanoDSF thermal shift assays of LptB2FG and assembled LptB2FGC (resolubilized from liposomes) in DDM micelles. Curves show the first derivative of the 350 nm fluorescence derived from a triplicate of experiments. LptB2FG is stabilized by 11 K upon addition of LptC (top and bottom) and by 9 K upon addition of LPS (top). LptB2FGC is stabilized by 2 K upon addition of LPS (bottom). (**E**) LILBID-MS spectra of LptB2FGC in DDM micelles in the apo state (light blue), after addition of 2 mole: 1 mole LPS: LptB2FGC (dark blue) and after addition of 2 mM ATP and 5 mM MgCl2 (purple) in the respective order. Highlighted regions show the fully assembled transporter with none, one or two bound LPS molecules. LPS is highlighted by a star in the cartoon.

For further analysis and confirmation of the complex formation, we re-solubilized the LptB_2_FGC-containing liposomes in detergent (DDM). Size-exclusion chromatography revealed a monodisperse peak with a retention volume 0.6 ml lower than that to LptB_2_FG alone, consistent with the increased molecular size of the LptB_2_FGC complex (Fig 2C). Our complex formation data are further supported by SDS-PAGE and blue-native PAGE (Fig S3A, C). Interestingly, thermal shift assay data of the protein in detergent micelles revealed an 11 K increase of the melting temperature in the presence of LptC (Fig. 2D), demonstrating a substantial stabilization of the complex through LptC binding. We then investigated the influence of LPS on the melting temperature of the individual proteins. LPS had no observable impact on the melting temperature of LptC alone (Fig S1). Surprisingly, addition of LPS to LptB_2_FG causes a similar shift in melting temperature of approx. 9 K. This finding is particularly notable given that LPS stimulates ATPase activity, whereas LptC reduces it. Addition of LPS to LptB_2_FGC only results in a modest 2 K increase in melting temperature compared to the apo LptB_2_FG complex. These results suggest that both LptC and LPS independently contribute to the stabilization of the ABC transporter.

We then probed LptB_2_FGC re-solubilised in DDM micelles with Laser-Induced Liquid Bead Ion Desorption Mass Spectrometry (LILBID-MS), to further investigate stoichiometry and functionality of the complex. The apo state of LptB_2_FGC yields a well-resolved spectrum in which we were able to assign almost every mass (Fig S2). We observed a dominant species of 158 kDa corresponding to the mass of the LptB_2_FGC complex in the expected 2:1:1:1 stoichiometry. A part of LptB_2_FGC complexes seems to be associated to LPS as we also observe a species of 162 kDa. Upon addition of a two-fold excess of LPS to the LptB₂FGC complex, we observed a pronounced increase of the population corresponding to the LPS-bound complex, accompanied by a decrease in the relative intensity of species representing smaller LptB₂FGC subcomplexes (Fig. 2E). Additionally, a new population corresponding to a complex with two bound LPS molecules emerged. The increased peak intensity can be interpreted as enhanced complex stability, as the same laser intensity results in reduced dissociation of the complex. We speculate that in addition to LPS binding in the proposed binding pocket of the TMDs, the second LPS could bind to either the βJR of LptF, LptC or in between. Treatment of the LptB₂FGC complex with ATP and magnesium chloride resulted in the dissociation of most bound LPS, as evidenced by a shift in mass peaks toward lower molecular weights corresponding to the unbound complex. This observation indicates that LptB₂FGC actively extrudes LPS into solution. Notably, LPS dissociation was accompanied by a further decrease in peak intensity, suggesting a reduction in complex stability following substrate release.

### Increased dynamics in the TMH of LptC during LPS transport

The available crystal and cryo-EM structures of LptB₂FG and LptB₂FGC were determined in detergent micelles or lipid nanodiscs, environments that do not fully replicate the native membrane context in which LPS extraction occurs (*15, 16, 21*). To investigate LPS transport by LptB₂FGC under more physiologically relevant conditions, we employed solid-state NMR (ssNMR) spectroscopy to study the complex reconstituted in liposomes. First, we recorded spectra of just LptC reconstituted in a POPE/POPG lipid bilayer. Previously, the chemical shifts of LptC were fully assigned using a soluble construct lacking the transmembrane helix (*18*). To reduce spectral overlap and to obtain site-specific insights into LptC within the full complex, we selectively isotopically labeled all lysine residues. Lysines are well distributed throughout the LptC primary sequence (Fig. 3A) and, importantly, do not undergo isotope scrambling, making them reliable reporters for ssNMR analysis (*38*). A similar labeling strategy has previously been employed in a study of the ABC transporter MsbA (*39*). To begin, we recorded an hNCA spectrum of uniformly labeled^15^N,^13^C-Lys-LptC, which exhibited a well-dispersed signal pattern (Fig. 3B). Labeling the 12 lysine residues resulted in 8 distinct signals in the spectrum. Of these, 6 resonances were assigned by transferring previously established solution NMR assignments (Fig. S4). To complete the assignment and validate the transferred resonances, we generated a series of lysine-to-alanine mutants, which also served as controls (Fig. S5). All observable βJR resonances display highly similar ^13^C and ^15^N chemical shifts, indicating that the conformation of the βJR domain in LptC remains largely unchanged in the presence of linker and TMH and when the TMH is embedded in a lipid environment. In contrast, the signals corresponding to the TMH residues K3 and K27 are significantly broadened and exhibit reduced intensity compared to the sharper, more intense signals from βJR lysines (K88, 90, 97, 100, 102, 162, 170) (Fig 3B). This broadening most likely results from conformational heterogeneity of K3 and K27 in the LptC_TMH_ as they are structurally less constraint than residues in the βJR.

**Figure 3:**
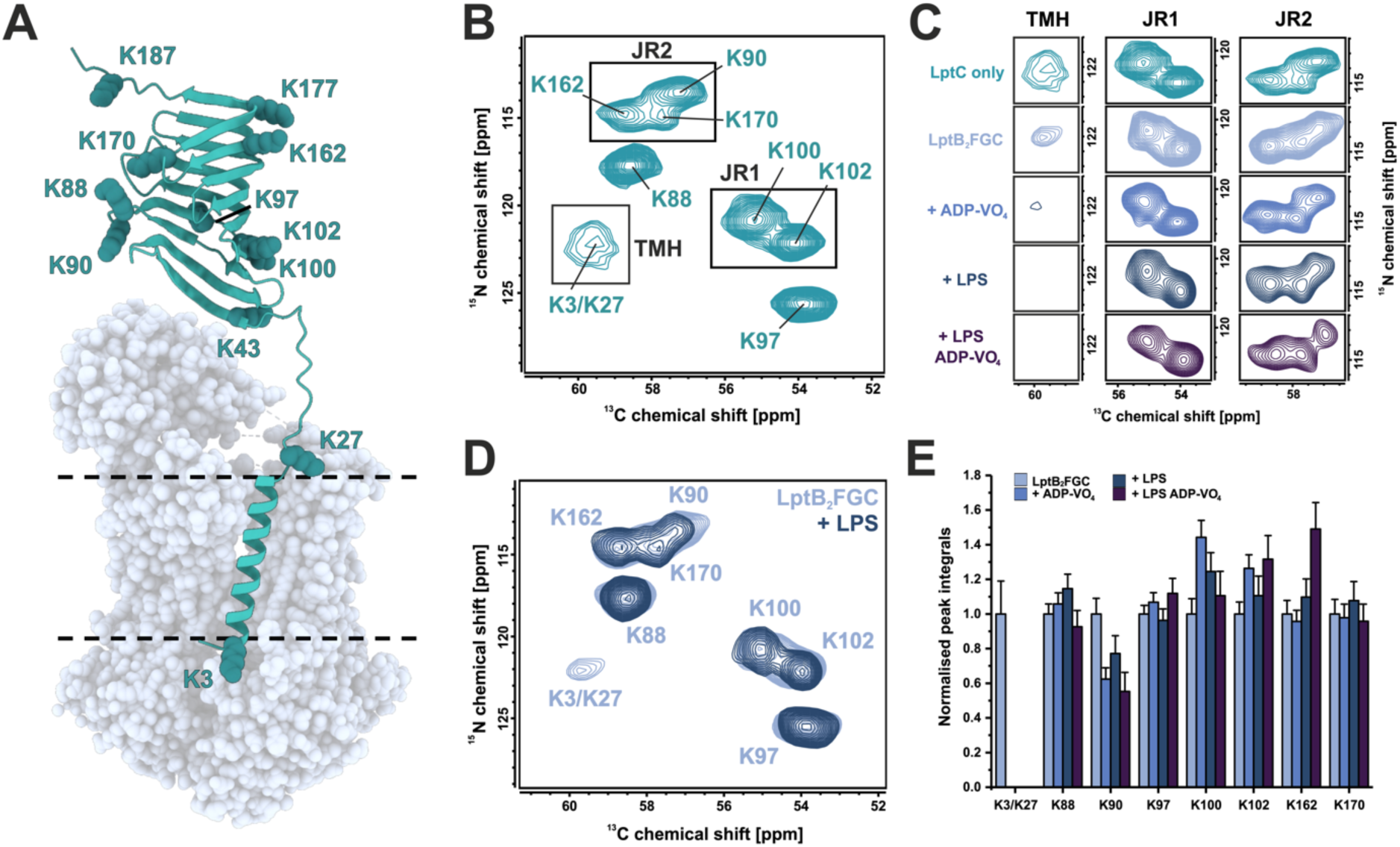
Probing LptC dynamics in the translocation process of LptB2FGC. **(A)** Alpha fold structure of *E. coli* LptC (AF-A0A0H3GZ01-F1-v4) with all lysines shown as spheres. The structure of LptB2FG (adapted PDB 6MJP) is shown as transparent spheres to emphasize the LptB2FGC complex. **(B)** 2D ^13^C, ^15^N hNCA of [^13^C, ^15^N-K]-LptC in liposomes under 14 kHz magic angle sample spinning at an 850 MHz NMR spectrometer. Spectral regions of interest are labeled as transmembrane helix (TMH), β-jellyroll region 1 (JR1) and β-jellyroll region 2 (JR2). **(C)** Selected spectral regions (TMH, JR1 and JR2) in different states of the LPS translocation cycle and **(D)** comparison of the full spectrum of LptB2FGC ± LPS. The TMH signal is significantly reduced upon LPS-binding, ADP.VO4-trapping or both. LptB2FGC was prepared by co-reconstitution of LptC and LptB2FG at 1:1 mole/mole in POPE/POPG. LPS was incorporated by using 2 mole LPS per mole LptB2FGC. **(E)** Integrals corresponding to the volume of 2D resonances from lysines of LptC derived from the spectra of C. The integrals were normalized twice, first to the sum of all lysine integrals to correct the difference in sample amount and second to the integral of each peak of the apo LptB2FGC sample. The error bars represent the noise that contributes to the signal integral determined by the root mean square of noise-areas from the respective 2D spectra.

Next, we studied ^15^N,^13^C-Lys-LptC within the LptB_2_FGC complex. To ensure that only signals from the bound form of LptC were observed in the hNCA spectra, we co-reconstituted LptB_2_FG and ^15^N,^13^C-Lys-LptC in a 1:1 molar ratio, thereby minimizing contributions from unbound LptC. The chemical shifts of the LptC βJR signals remained largely unchanged; however, variations in integral signal intensities were observed, which may reflect weak or transient interactions between the βJR of LptC and LptF (Fig. S6A). Interestingly, the transmembrane signals corresponding to K3 and K27 appeared sharper in the context of the LptB₂FGC complex compared to LptC alone, potentially indicating a restriction in the conformational flexibility of the LptC_TMH_ upon complex formation (Fig. 3C). We next prepared samples of LptB_2_FGC in the vanadate-trapped, LPS-bound and in the ADP-VO_4_-trapped-LPS-bound state. Successful trapping of the high-energy states was confirmed by ^31^P NMR (Fig. S6B). No significant chemical shift changes across these conditions were observed for the βJR domain. However, in all these states, the TMH signal from K3/K27 was markedly reduced or completely undetectable (Fig. 3C and Fig S6C). Full dissociation of the LptC from the complex should result in a TMH signal similar to the one observed for membrane-bound LptC. A loss of signal in the hNCA spectrum can be caused by enhanced molecular motions resulting in an insufficient cross polarization transfer and/or conformational exchange on the intermediate NMR timescale. The data therefore demonstrate an increase in dynamics of LptC_TMH_ within the transporter complex upon LPS binding, ADP.VO_4_-trapping or both. This observation indicates an allosteric modulation/destabilization of the LptC_TMH_ interaction with LptB_2_FG, mediated by the NBDs and/or LPS.

In comparison to LptC_TMH,_ the changes of lysines peaks in the βJR of LptC are only subtle. We do not observe any major changes in the chemical shift in any of the states. The LptC_βJR_ does not appear to undergo signficant conformational changes during the translocation cycle of LptB_2_FG, at least in the absence of LptA. However, we observe notable changes in the peak integrals of several lysine residues (Fig 3D). For comparison, peak integrals were normalized to the total signal intensity of all lysine peaks in the spectrum. K90 and K162 seem to be affected most in the states of LPS translocation with K90 displaying decreased and K162 increased signal intensity. K90 is the closest visible lysine to the expected interaction interface with LptF (Fig 3A), suggesting that its altered intensity may reflect dynamic interactions at this site during the transport cycle. The interaction between of the LptC_βJR_ and the LptF_βJR_ has been described to be highly dynamic in the LPS bound cryo-EM structures (*16*). Our data would support a transient interaction of the LptC-LptF βJR domains, which is altered in the presence of LPS, ADP-VO_4_ or both.

### DNP enhanced TEDOR reveals two distinct states of LptC during LPS translocation

The observed changes in the dynamics of LptC_TMH_ in LptB_2_FGC raise the question of whether LptC_TMH_ remains deeply inserted in the transporter throughout the different catalytic states. To address this question, we probed intermolecular contact between LptC and LptB_2_FG by preparing a mixed-labelled complex from ^13^C-LptB_2_FG and ^15^N-LptC (Fig. 4A). Through-space ^13^C-^15^N dipole-dipole couplings can then be visualized by DNP-enhanced ^13^C-^15^N-TEDOR experiments. The signal enhancement provided by DNP enables the detection of weaker spin couplings, while the cryogenic conditions required for DNP (100 K) effectively freeze molecular motions, yielding spectra that capture all coexisting conformational states of the transporter. The TEDOR mixing time of 6.5 ms was selected to optimize the detection of long-range intermolecular contacts. The ^13^C-^15^N-TEDOR spectrum obtained for apo state ^15^N-LptC-^13^C-LptB_2_FG shows several signals. Most of them result from natural abundance cross peaks from N-CO, N-CA and N-CX coupling due to 1.1% ^13^C natural abundance in ^15^N-LptC and 0.4% ^15^N natural abundance in ^13^C-LptB_2_FG (Fig. 4B, for a detailed discussion of these contributions see reference (*40*)). The origin of these cross peaks has been verified in control spectra recorded on LptC-^13^C-LptB_2_FG and ^15^N-LptC-LptB_2_FG (Figs. 4 D, E).

**Figure 4:**
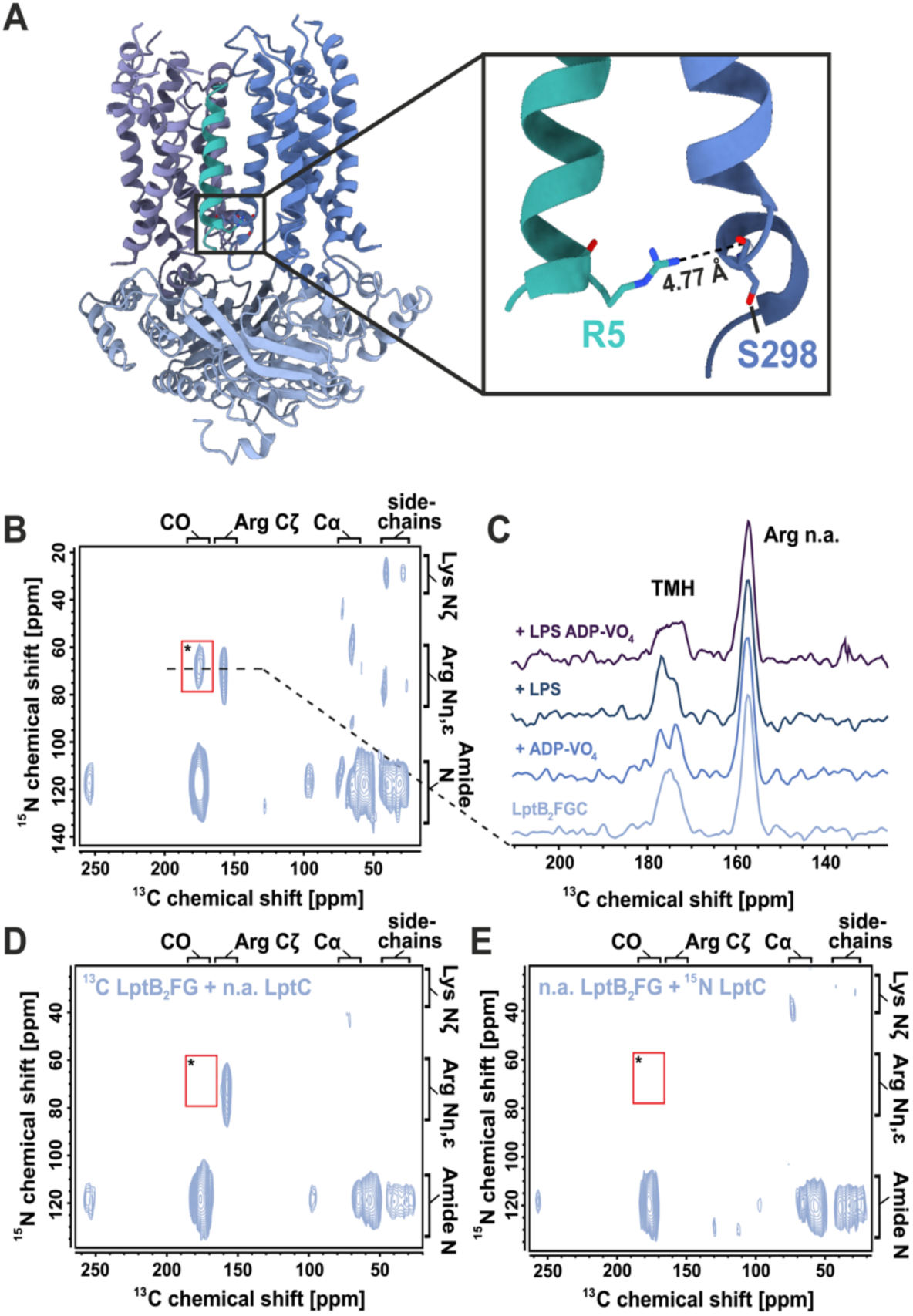
The conformational landscape of the LptCTMH probed with DNP enhanced solid-state NMR on the mixed-labelled ^15^N-LptC-^13^C-LptB2FG complex. **(A)** *E. coli* LptB2FGC cryo-EM structure in the LPS-bound nucleotide free state (PDB: 6MI7). Zoom-in on the assigned cross-contact between Arg-5 Nη,ε of LptC and the backbone C’ of Ser-298 of LptF. **(B)** DNP-enhanced ^13^C, ^15^N-TEDOR spectra of ^15^N-LptC-^13^C-LptB2FG complex in liposomes. The specific LptC-LtB2FG contact between Arg5 and Ser298 is labelled by an asterisk. All other cross peaks arise from natural abundance correlations (see D) and (E)) **(C)** 1D ^13^C slices through peak * along ω1(^15^N) = 70 ppm for different states of LPS translocation. Spectra are normalized on the Cζ arginine natural abundance signal (157 ppm). The corresponding 2D TEDOR spectra are shown on Fig. S8. **(D)** Natural abundance control spectra of LptC-^13^C-LptB2F and **(E)** ^15^N-LptC-LptB2F in liposomes at 100 K. Here, all cross peaks arise from natural abundance correlation of 0.4% ^15^N with 100% ^13^C in (D) and 1.1 % ^13^C with 100 % ^15^N in (E). The intermolecular cross-contact * cannot be detected with these labelling schemes. All DNP TEDOR spectra were recorded with a mixing time of 6.5 ms at 100 K.

**Figure 5:**
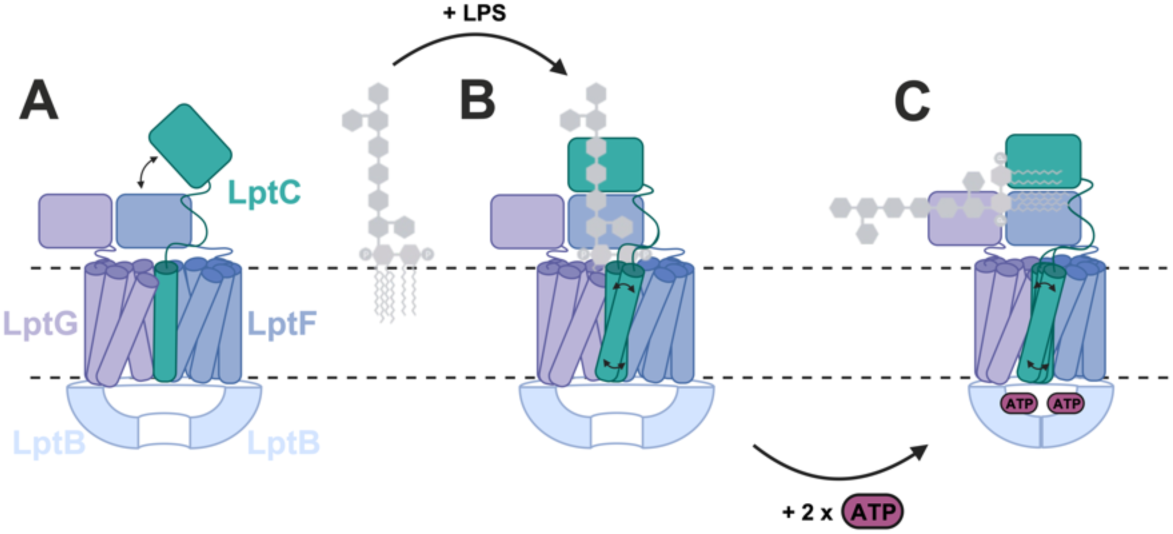
Proposed model for the role of LptC in LPS translocation by LptB2FGC. **(A)** In the apo state (absence of both LPS and nucleotide), LptCTMH adopts a conformation in which it is conformation sandwiched between LptF and G. The βJR of LptF and LptC engage in transient interactions. **(B)** Upon LPS binding to the nucleotide-free state, LptCTMH undergoes increased conformational exchange, switching between two distinct LptB2FG-bound states, indicative of enhanced helix dynamics. **(C)** In the Vanadate-trapped, LPS-bound state, LptCTMH exhibits further increased dynamics, accompanied by structural rearrangements in the LptCβJR domains due to its association with LptF. The figure was created using BioRender (*52*).

Importantly, cross peak * (μ_2_(^13^C) = 70 ppm, μ_1_(^15^N) = 175 ppm) only occurs in the mixed-labeled complex and must therefore arise from an intermolecular contact between LptC and LptB_2_FG (Fig. 4A, B). The nitrogen chemical shift of the observed inter molecular contact is unique for the Nη,ε of an arginine residue whereas the ^13^C chemical shift at 175 ppm originates from either a carbonyl backbone signal (C’) or a sidechain Cψ/8 carbon of E, Q, D or N. To identify potential candidates for the observed intermolecular contact, we analyzed the distances between the Nη,ε atoms of LptC arginine residues and either side chain (Cψ/8) or backbone (C’) atoms of LptB₂FG, based on available cryo-EM and crystal structures of LptB₂FGC (PDB IDs: 6MJP and 6MI7). The analysis revealed only a few potential contact candidates. Of these, only one falls within the TEDOR detection range (<5 Å): the Nη,ε of R5 from LptC_TMH_ and the backbone C’ of S298 in LptF (4.8 Å). To confirm that the observed cross peak originates from the R5-LptC_TMH_ – S298-LptF interaction, we introduced a single-point mutation (R5L) in LptC. This mutation abolished the corresponding intermolecular cross peak (Fig. S7), supporting a definitive assignment to the specific LptC– LptF interface. The R5-LptC_TMH_ – S298-LptF contact thus serves as reporter for monitoring changes of the LptC_TMH_ interaction during different states of LPS translocation.

We next repeated the TEDOR experiment for the ADP-VO_4_-trapped state, the LPS-bound state and the combined ADP-VO_4_-trapped/LPS bound state (Figs. 4C, S8). In three conditions, the intermolecular cross peak (*) between LptC_TMH_ and LptF was clearly observed. This was unexpected, as previous studies have suggested that LptC_TMH_ dissociates from the transporter in the presence of either LPS or ADP-VO_4_ (*16, 27*). In particular, cryo-EM data of LptB_2_FGC in the vanadate-trapped state show insufficient space for the LptC_TMH_ to remain positioned between LptG and LptF (*16*). Our results, however, demonstrate that at least the lower part of the helix remains in contact with LptF throughout the translocation process. Interestingly, the line shape of the cross peak - which reflects the available conformational space sampled at the site - varies across the different functional states (*41*). In both the ADP-VO₄–trapped and LPS-bound states, two distinct signals are observed, indicating that LptC does not adopt a single, rigid conformation. In the combined LPS-bound / vanadate-trapped state, the cross peak becomes notably broader and decreases in intensity, consistent with increased conformational heterogeneity. Taken together, these observations suggest that the presence of LPS and vanadate-trapping induce a conformational exchange between two distinct states on the microsecond timescale. Importantly, in both states, LptC_TMH_ remains in contact with the LptB_2_FG complex. In the combined LPS/vanadate-trapped state, LptC_TMH_ appears more heterogenous and to sample multiple conformations between these two states, leading to a broader signal in the TEDOR spectra.

## Discussion

ABC transporters play a critical role in mediating substrate transport across bacterial membranes. However, LptB₂FGC is unique among known ABC transporters in that it incorporates an associated bitopic membrane protein, LptC, as part of its architecture (*22, 42*). Despite all the structural and functional data, the role of LptC remains elusive – particularly the contribution of its transmembrane helix (TMH) to LPS extrusion. Here we present a possible role for the LptC_TMH_ as a mechanical transducer facilitating coordination between ATP hydrolysis and LPS binding and transport.

Our findings demonstrate that both LptB_2_FG and LptB_2_FGC exhibit an LPS-dependent stimulation of ATPase activity *in vitro* (Fig. 2B.) LptC association reduces the absolute activity even in the presence of LPS, suggesting that LptC counteracts LPS stimulation. Increasing amounts of LptC further reduce the basal ATPase activity to a minimum of 45 % in the apo state (Fig. 2A). This effect may serve to minimize ATP consumption in the absence of substrate. As LptC and LptF/LptG are encoded under different operons (*43*), the organism could be able to fine-tune the ATPase activity of LptB_2_FG by changing the expression level of LptC.

The influence of LptC on ATPase activity is particularly interesting, when comparing the stability of LptB_2_FG and LptB_2_FGC. We observed strong thermal stabilization of LptB_2_FG in micelles upon binding to LptC, LPS or both (Fig. 2D). LptC has previously been shown to reduce conformational heterogeneity of LptB_2_FG in a membrane environment (*44*), supporting a model in which LptC stabilizes the transporter in an inactive or primed state until LPS is available. Upon LPS binding, a marked increase in thermal stability of LptB_2_FGC is observed in micelles. LILBID-MS spectra revealed the presence of two bound LPS molecules (Fig. 2E). This suggests that this stabilization arises not only from LPS occupying the hydrophobic cavity within the transmembrane domains (TMDs), but also from interactions with the LPS binding sites in the β-jellyroll (βJR) domains. These in vitro observations are consistent with in vivo findings that LPS binding stabilizes the Lpt bridge complex and extends its lifetime (*12*).

Tracking the position and dynamics of LptC_TMH_ during LPS transport remains challenging, as currently available structural data fail to resolve the helix in the AMP-PNP and ADP-VO_4_-trapped states (*15, 16*). To overcome this limitation, we employed both ambient and DNP-enhanced cryogenic solid-state NMR to monitor changes in helix dynamics and interactions under native-like membrane conditions (Figs. 3, 4). Our results reveal that LPS and nucleotide binding lead to increased conformational dynamics of LptC_TMH_. The helix undergoes conformational exchange between an associated state—where it is positioned between TM1 of LptG and TM5 of LptF—and a second, more dynamic state, which we propose is oriented more toward the exterior of the proposed binding pocket. We speculate that this alternative binding site of the TMH is occupied by a POPG lipid, as observed in the LptB_2_FGC cryo-EM structure (PDB:6MI7 (*16*), Fig S9). In this alternative site, the LptC_TMH_ remains in close proximity to LptF but adopts a distinct conformation within the transporter, which is consistent with our TEDOR data (Fig. 4).

Recently, *in vivo* data proposed a mechanism in which the LptC_TMH_ is displaced upon LPS as well as nucleotide-binding (*27*). However, our findings indicate that LptC_TMH_ does not fully dissociate from the complex during LPS transport. Instead, LPS binding induces conformational exchange rather than complete displacement of the helix. This dynamic behavior provides a mechanistic explanation for the dual regulation of ATPase activity – where LPS stimulates, but LptC simultaneously inhibits the transporter. Moreover, structural data suggested that full insertion of the LptC_TMH_ may interfere with substrate transport as the ADP-VO_4_-trapped conformation lacks sufficient space to accommodate LptC_TMH_ (*16*). These findings support a model in which LptC_TMH_ functions as a mechanical transducer, modulating ATP hydrolysis efficiency in response to substrate availability. Thus, LptC_TMH_ may sterically hinder the collapse of the LptFG transmembrane domains (TMDs); however, in the presence of LPS, its increased dynamics render the TMD collapse more favorable, thereby enhancing ATPase activity. In this way, LptB_2_FGC may enable an efficient ATP consumption in proportion to the amount of transported LPS molecules.

Very recent EPR distance measurements between spin-labelled positions within LptC_TMH_ and LptF/G in the full Lpt transporter complex, conducted in the presence of LPS with and without ATP, revealed a ∼10 Å increase in distance between labels, suggesting a significant positional shift of the helix (*45*). However, data for the LPS-free apo state were not reported. This observation appears at first glance to contradict our DNP TEDOR results, which demonstrate a clear contact between LptC and LptF under comparable conditions (Fig. 3C), albeit with notable differences in structural heterogeneity. Importantly, the increase in EPR-measured distance does not necessarily imply a lateral displacement consistent with domain dissociation. Instead, it may reflect a conformational rearrangement involving the whole complex in which the LptC_TMH_ helix toggles between two positions as part of the Lpt extrusion cycle as suggested above.

The association and interactions of the β-jelly roll (βJR) domains among the various components of the Lpt system appear to be essential for forming the complete LPS bridge; however, these interactions are transient and can be modulated by LPS or ADP-VO₄ binding (*16*). A comprehensive in vitro assessment of substrate-dependent bridge formation will clearly require the inclusion of LptA and LptDE to fully reconstitute the periplasmic Lpt bridge. Finally, the observed conformational flexibility of LptC_TMH_ leads us to speculate that the mobility of this helix may also be critical for providing the spatial accommodation necessary for LptF–C–A βJR association.

In combining all of our data, we propose a model for the role of LptC_TMH_ for LPS translocation of LptB_2_FGC, which includes the dynamic information from our NMR studies (Fig 6). LptC_TMH_ is bound to the TMD of LptB_2_FG and adopts one preferred conformation as shown by the signals of LptC-K3 and K27 (Fig. 3) and the contact between LptC-R5 and LptF-S298 (Fig. 4). Upon LPS binding, the mobility of LptC_TMH_ is enhanced as deduced from the observed signal loss of LptC-K3 and K27, while the LptC-R5 - LptF-S298 contact remains and is splitting into two signals. In this state, the LptC_TMH_ appears to be exchanging between positions inside and outside of the binding pocket. Such increased mobility leads to reduced steric hindrance, which then supports collapse of the binding pocket during LPS translocation. In the vanadate--trapped state, LptB_2_FG like all ABC transporters adapt a more rigid conformation, but LptC_TMH_ with and without LPS remains mobile for an easier re-association with LptB2FG in the apo state. The βJR association of LptC with LptF is modulated upon LPS and ADP-VO_4_ binding We speculate that the βJR bridge formation of LptF and LptC is enhanced in the presence of its native substrate. However, the observed changes are subtle, and a comprehensive assessment will require investigation of additional components of the Lpt system, such as LptA, which may influence bridge formation and stability. Overall, our findings provide new insights into LptC-dependent regulation of the LptB₂FG complex and lay the groundwork for the development of therapeutic strategies targeting regulatory elements within LptB₂FGC.

## Materials and Methods

### Bacterial Strains, Plasmids and cloning of Lpt genes

*E. coli* strain BW25113 (a derivative of *E. coli* K12 strain) was kindly provided by Prof. Klaas Martinus Pos (Goethe University Frankfurt). *E. coli* strain C43 (DE3) was kindly provided by Prof. Hartmut Michel (Max Planck Institute of Biophysics, Frankfurt). *E. coli* strains BL21 (DE3) and DH5alpha were purchased from Thermo Fisher Scientific. The genes encoding the LptB_2_FG complex were cloned into the pETDuet-1 vector, with LptB (bearing a N-terminal His_6_-tag) inserted into the first multiple cloning site, and LptF and LptG cloned into the second site, resulting in the expression plasmid pET-Duet1-LptB_2_FG. The gene encoding full-length LptC, including its transmembrane helix, was cloned into pBAD33.1 vector with a C-terminal His_6_ tag.

### Site-directed mutagenesis

Single lysine mutations in LptC were introduced using the plasmid pBAD33.1-LptC-His_6_ as template. Mutation-specific primers were designed using the QuickChange Primer Design Tool (Agilent). The sequences of forward and reverse primers of each mutation are listed in Table S1. PCR conditions were optimized individually for each mutant, and successful mutagenesis was confirmed by DNA sequencing (Microsynth Seqlab GmbH). All LptC mutants were expressed, purified, and reconstituted following the same protocol as for wild-type LptC, as described below.

### Expression and purification of LptC

A total of 100 ng of plasmid DNA was transformed into *E. coli* BL21(DE3) cells and plated on LB agar containing chloramphenicol for selection. A single colony was picked and inoculated into 100 mL of LB medium supplemented with chloramphenicol, followed by overnight incubation at 37 °C with shaking at 190 rpm. Cells from a 5 mL aliquot of the overnight culture were harvested, washed with fresh LB medium, and used to inoculate 500 mL of M9 minimal medium to initiate protein expression.

Cells were further grown overnight at 37 °C with shaking at 190 rpm in M9 minimal medium. For protein expression, cells from the overnight M9 culture were harvested by centrifugation and resuspended in fresh M9 medium supplemented with amino acids, adjusted to a starting OD_600_ of 0.2.

For selective amino acid labeling, the M9 medium (composition: 3.39 g K₂HPO₄, 1.5 g KH₂PO₄, 0.25 g NaCl, 0.5 g NH₄Cl, 2 g glycerol, 1.0 mL of 1 M MgSO₄, and 0.5 mL of 0.01 M FeCl₂ per 500 mL culture) was supplemented with ¹³C,¹⁵N-labeled lysine (Silantes, Munich, Germany), while all other amino acids remained unlabeled.

For uniform nitrogen isotope labeling, ¹⁵N-labeled NH₄Cl was used as the sole nitrogen source in the M9 medium, without supplementation of any amino acids.

After the media exchange, cells were incubated at 37 °C with shaking at 190 rpm for 2 hours until the OD_600_ reached 0.6–0.7. The temperature was then reduced to 20 °C, and the cells were allowed to adapt for 1 hour. Protein expression was subsequently induced with 0.2% (w/v) arabinose and carried out at 20 °C and 190 rpm. After 18 hours of expression, cells were harvested by centrifugation and resuspended in lysis buffer (50 mM HEPES, 300 mM NaCl, 5% glycerol, 5 mM MgCl₂, pH 7.5) supplemented with protease inhibitors and DNase I.

Membranes were prepared by disrupting resuspended cells using a high-pressure cell disrupter at 1.8 kbar (three passes), followed by centrifugation at 4,000 × *g* (7,000 rpm, rotor F0850) for 15 minutes to remove cell debris. The membrane was pelleted by and an ultracentrifugation step at 223,000 × *g* (55,000 rpm, rotor 70 Ti) for 1 hour at 4 °C.

The membrane pellet was solubilized in buffer A (50 mM HEPES, 300 mM NaCl, 5% glycerol, pH 7.5) containing 1.0% lauryldimethylamine-N-oxide (LS) and a complete protease inhibitor tablet. Solubilization was carried out at 4 °C for 1 hour. Insoluble material was removed by ultracentrifugation at 223,000 × *g* for 40 minutes (same rotor and temperature).

The resulting supernatant was applied to Ni-NTA beads (Qiagen, Hilden, Germany) pre-equilibrated with buffer A containing 5 mM imidazole. After 1 hour of binding at 4 °C, the protein was eluted with buffer A containing 0.025% LS and 350 mM imidazole.

The eluate was concentrated using a 10 kDa molecular weight cutoff Amicon concentrator, and imidazole was removed via a PD-10 desalting column. Protein concentration was determined using a Nanodrop spectrophotometer, using an extinction coefficient of 29,910 M⁻¹cm⁻¹ and a molecular weight of 22.8 kDa. Protein purity, homogeneity, and monodispersity were assessed by SDS-PAGE and size exclusion chromatography using a Superdex 200 Increase 10/300 GL column (Fig. S3).

### Expression and purification of LptB_2_FG

Protein expression and purification were performed as previously described, with minor modifications. Membranes were solubilized in lysis buffer (50 mM HEPES, 300 mM NaCl, 5% glycerol, pH 7.5) supplemented with 1% n-dodecyl-β-D-maltoside (DDM), 2 mM ATP, and a complete protease inhibitor tablet. Solubilization was carried out at 4 °C with shaking for 1 hour. Insoluble material was removed by ultracentrifugation at 223,000 × *g* (55,000 rpm, rotor 70 Ti) for 40 minutes at 4 °C.

The remaining supernatant was applied to Ni-NTA beads (Qiagen, Hilden, Germany) pre-equilibrated with buffer containing 5 mM imidazole. After 1 hour of binding at 4 °C, the resin was washed sequentially with buffer containing 15 mM and 20 mM imidazole to remove non-specifically bound proteins. The target protein was then eluted using buffer containing 200 mM imidazole.

### Thermal shift assays by NanoDSF

Purified LptC, LptB_2_FG and re-solubilised LptB_2_FGC in detergent containing purification buffer (50 mM HEPES pH 7.5, 150 mM NaCl, 5 mM MgCl_2_, 5 % glycerol, 0.05 % DDM). Samples were incubated for 30 minutes at room temperature either in the absence or presence of a 2-fold molar excess of Ra-LPS (Sigma-Aldrich Chemie GmbH, Taufkirchen, Germany). Thermal unfolding profiles were measured using a Prometheus Panta (NanoTemper Technologies, Munich, Germany). A temperature gradient of 1 °C/min from 25 °C to 85 °C was applied. Intrinsic tryptophan fluorescence was monitored at 330 nm and 350 nm. Data analysis was performed using the manufacturer’s software, and thermal transitions were identified by plotting the first derivative of the 350 nm fluorescence intensity curves.

### Laser induced Liquid ion bead desorption (LILBID) mass spectrometry

Proteins solubilized in detergent were buffer-exchanged into 20 mM Tris-HCl (pH 7.5) containing 0.05% DDM prior to analysis by laser-induced liquid bead ion desorption mass spectrometry (LILBID-MS). For LPS binding studies, Ra-LPS was added at a 2:1 molar ratio (LPS:protein) following buffer exchange. Samples were incubated for 10 minutes at room temperature. To study LPS dissociation, 2 mM ATP and 5 mM MgCl₂ were added to the LPS-bound protein complexes, followed by an additional 10-minute incubation at room temperature.

For MS analysis, 4 µL of each sample were loaded into a piezo-driven droplet generator (MD-K-130, Microdrop Technologies GmbH, Germany), generating aqueous droplets of approximately 50 µm diameter at a frequency of 10 Hz. Upon introduction into high vacuum, droplets were irradiated with an IR laser (λ = 2.8 µm, maximum energy output of 23 mJ per pulse).

Free ions generated upon desorption were accelerated and transferred into a custom-built time-of-flight (TOF) mass analyzer. For this, the voltage between the first and the second lens in the Wiley McLaren type ion optics was set to −4.0 kV while the first lens was pulsed to −6.6 kV 5-25 µs after the irradiation for a time of 370 µs. Detection was carried out using a custom Daly-type detector.

### Reconstitution of the Lpt proteins

A lipid mixture of POPE:POPG (4:1 molar ratio) (Avanti Polar Lipids, Inc.) was used for protein reconstitution at a lipid-to-protein molar ratio of 100:1. Lipids were initially dissolved in a chloroform:methanol (2:1, v/v) mixture and dried under a gentle stream of nitrogen gas. Residual solvent was removed by vacuum rotary evaporation.

The resulting dried lipid film was rehydrated in 50 mM HEPES buffer (pH 7.5) and subjected to three freeze–thaw cycles, alternating between liquid nitrogen and a 37 °C water bath. The lipid suspension was then extruded 11–13 times through polycarbonate membranes with a 0.2 µm pore size to generate unilamellar vesicles. Liposomes were destabilized with 3 mM DDM for LPTB_2_FG and 2mM LS for LptC according to previously used reconstitution protocols (*46*). Ra-LPS was incorporated into the liposomes as described before (*13*). Sample homogeneity was assessed using a discontinuous sucrose gradient prepared by sequentially layering 1 mL each of 10%, 30%, 50%, and 70% (w/v) sucrose in 50 mM HEPES buffer (pH 7.5). Approximately 300 µL of the sample was carefully layered on top of the gradient and centrifuged at 28,000 rpm for 16 hours at 4 °C using a Beckman 50.1 Ti rotor.

### Co-reconstitution and re-solubilisation of the LptB_2_FGC

Purified LptB₂FG and LptC were mixed and incubated for 5 minutes at room temperature prior to addition to destabilized liposomes. Reconstitution was followed by solubilization of the resulting proteoliposomes through the addition of 0.5% DDM in purification buffer and incubation with shaking at 4 °C for 1 hour.

Following solubilization, the sample was centrifuged at 200,000 × g for 1 hour at 4 °C. LptB₂FGC was further purified by size exclusion chromatography on a Superdex 200 Increase 10/300 GL column (see Fig. 2C).

### ATPase activity assay

The activity of LptB₂FG and LptB₂FGC was determined by measuring the release of inorganic phosphate (Pi) resulting from ATP hydrolysis, using the Malachite Green assay (*47*). Reaction mixtures contained LptB₂FG and LptB₂FGC proteoliposomes and varying concentrations of ATP (0.0–5.0 mM) in assay buffer comprising 50 mM HEPES (pH 7.5), 50 mM NaCl, and 20 mM MgCl₂. Reactions were incubated at 37 °C for 1 hour and terminated by addition of ice-cold stop buffer containing 20 mM H₂SO₄. Malachite Green reagent was then added, and the absorbance corresponding to Pi release was measured spectrophotometrically. The effect of LptC on ATPase activity (Fig. 2A) was determined by co-reconstituting increasing amounts for LptC together with LptB_2_FG into POPE/POPG liposomes (LptC: LptB_2_FG 0 - 2 mol/mol). The influence of LPS on ATPase activity of LptB_2_FG and LptB_2_FGC (Fig. 2B) was determined by reconstituting LptB_2_FG or LptB_2_FGC into POPE/POPG liposomes with increasing amounts of RA-LPS (0 – 15 mol/mol RA-LPS/LptB_2_FG or LptB_2_FGC). For the latter, a two-fold excess of LptC over LptB_2_FG was used to ensure saturation of the full complex.

Absolute ATPase activities were determined as follows: 2.87 ± 0.12 mol Pi/mol LptB₂FG·min⁻¹ (defined as 100% activity in Fig. 2A), 3.70 ± 0.08 mol Pi/mol LptB₂FG·min⁻¹ (for 0 mol LPS in Fig. 2B), 0.53 ± 0.04 mol Pi/mol LptB₂FGC·min⁻¹ (for 0 mol LPS in Fig. 2B). The differences between the values for LptB_2_FG are due to batch-to-batch variations.

### Ambient temperature and DNP enhanced solid-state NMR spectroscopy

For NMR experiments conducted at ambient temperature, proteoliposomes were incubated with 1 mM Gd³⁺-DOTA for 15 minutes as previously described (*48*). Samples were then pelleted by centrifugation at 200,000 × g and packed into 3.2 mm MAS rotors. Sample quantities were as follows: 7 mg of wild-type LptC for the LptC-only sample, 5–7 mg of LptC mutants, 3–4 mg of LptC for samples co-reconstituted with LptB₂FG.

Vanadate-trapping of reconstituted LptB_2_FG or LptB_2_FGC complex was carried out essentially as described before (*49*). Briefly, 20–25 mg of reconstituted LptB₂FG or LptB₂FGC in 50 mM HEPES buffer (pH 7.5) was mixed with ATP, MgCl₂, and sodium orthovanadate at a molar ratio of 1:400:400:120 (protein:ATP:MgCl₂:vanadate). The mixture was subjected to three freeze–thaw cycles (liquid nitrogen and 37 °C water bath) to enhance trapping efficiency, followed by incubation at 37 °C for 30 minutes. Following incubation, the sample was pelleted and washed thoroughly with buffer containing 50 mM HEPES and 5 mM MgCl₂ to remove excess reagents.

For DNP, small pellets of reconstituted samples were incubated overnight at 4 °C with 20 mM AMUPol in a cryoprotectant buffer consisting of 10% H₂O, 30% D₈-glycerol, and 60% D₂O. After incubation, the supernatant was carefully removed, and the pellets were transferred into a 3.2 mm zirconium oxide MAS rotor by centrifugation.

NCA spectra were recorded on a Bruker 850 MHz Avance III spectrometer equipped with a 3.2 mm DVT-HCN E-free MAS probe, using a magic-angle spinning (MAS) rate of 14 kHz at a sample temperature of 270 K (corresponding to an actual sample temperature of 5–10 °C). For the N–C cross-polarization (CP) step, a mixing time of 3.5 ms was applied with continuous wave (CW) decoupling at 100 kHz. Each NCA spectrum was recorded with 3072 scans, 80 increments, and a recycle delay of 1 s.

¹³¹P CP MAS NMR spectra were acquired under the same conditions (14 kHz MAS, 270 K) using the same probe. A CP contact time of 3 ms was used, followed by high-power proton decoupling during acquisition. The DNP enhanced MAS NMR spectra were recorded using a Bruker DNP system consisting of a 400 MHz WB Avance NEO spectrometer (Frankfurt) or a Avance III spectrometer (University Darmstadt), a 263 GHz Gyrotron as a microwave source and a 3.2 mm HCN-DNP-MAS probe. DNP measurements were carried out at 100 K and a sample-spinning rate of 8 kHz. The ^13^C-^15^N TEDOR experiment was acquired with a 6.5 ms mixing time and 32 increments with 3072 scans each. Spectral widths were 40.65 kHz in ω2 and 8 kHz ω1. We observed a DNP enhancement ε between 17-25 in the measured samples.

For all NMR experiments, standard settings for cross-polarization (CP) and decoupling were used. A typical ¹H 90° pulse had a duration of 3 μs. CP contact times were selected between 0.8 and 1.5 ms, depending on the experiment. High-power proton decoupling in the range of 70–100 kHz was applied using the SPINAL-64 decoupling scheme (*50*).

All spectra were processed using Bruker TOPSPIN 4.2.0.

## Acknowledgments

This work was funded by the CRC 1507 ‘‘Membrane-associated protein assemblies, machineries and supercomplexes’’. The authors thank Melanie McDowell, MPI for Biophysics, for access to their NanoDSF device, Cédric Laguri for providing the solution NMR assignment of LptC and Ute Hellmich, University of Jena and Klaas Martinus Pos, Goethe University Frankfurt for helpful discussions. We further thank Klaas Martinus Pos for providing the *E. coli* strain used in this work. We thank Julian Fischer and Inna Gjergji for help with protein expression. We also express our gratitude to Torsten Gutmann, TU Darmstadt for help with the additional DNP measurements.

## Author contributions

AK carried out all NMR and biochemical experiments of LptB_2_FG and LptB_2_FGC. JK carried out all NMR and biochemical experiments of LptC. JK and SS performed cloning and established the initial protocol for the expression and purification of the used Lpt proteins. TR carried out all LILBID-MS experiments and analyzed the data together with NM. CG conceived the project. AK, JK, JB and CG designed experiments, analyzed and interpreted data. AK, JK and CG designed figures and wrote the paper. All authors contributed to review and editing of the manuscript.

## Competing interests

The authors declare no competing interests.

## Data and materials availability

All data needed to evaluate the conclusions in the paper are present in the paper and/or the Supplementary Materials.

## Notes

### Competing Interest Statement

The authors have declared no competing interest.

