## Supporting data for "Conformational plasticity of LptC regulates lipopolysaccharide transport by the LptB_2_FGC complex"

**This file includes:**

- Supplementary biochemical data
- Supplementary NMR spectra of LptC mutants
- $^{31}P$ NMR spectra of $ADP.VO_4$-trapped $LptB_2FGC$
- Supplementary DNP spectra

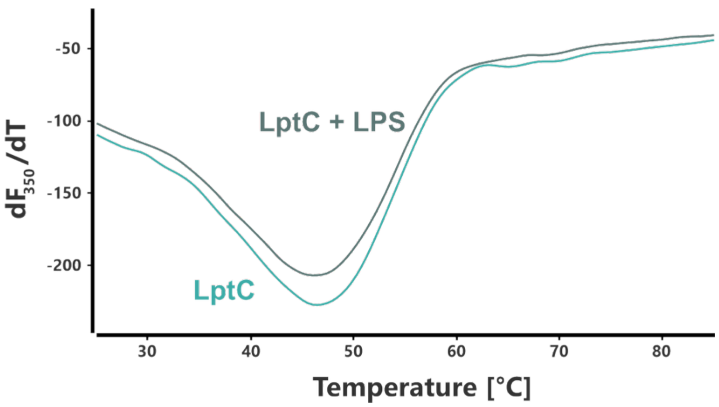


**Fig. S1:** NanoDSF thermal shift assays of LptC and LptC with LPS at 1:2 mole ratio in LS micelles. Curves are derived from a triplicate of experiments. No change in the melting temperature $T_m$ 46.52 ± 0.15 °C can be observed after the addition of a 2-fold access of LPS (46.18 ± 0.25 °C). For experimental details see corresponding paragraph in the materials and methods section.

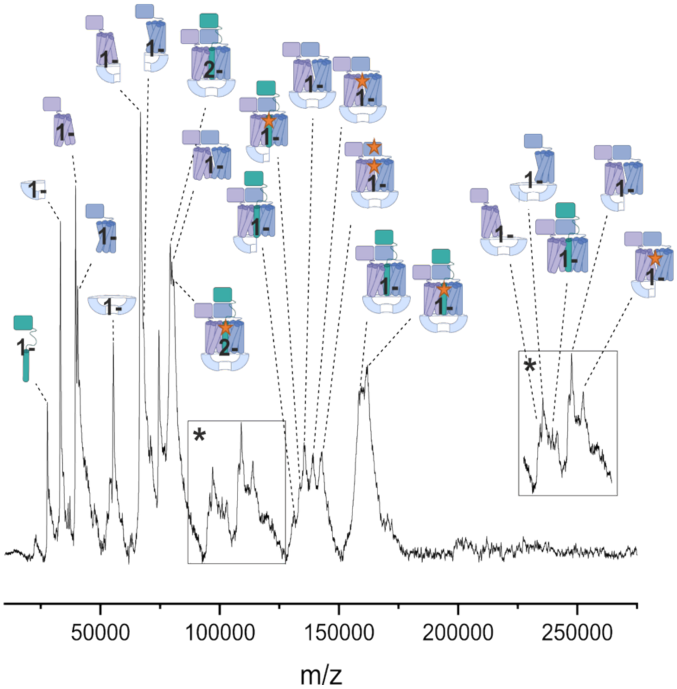


**Fig. S2:** LILBID-MS spectra of the $LptB_2FGC$ apo state in DDM micelles. Almost every signal could be assigned to complexes, subcomplexes and subunits of $LptB_2FGC$. Importantly dominant species of 158 kDa (singly and doubly charged) could be observed, originating from the assembled $LptB_2FGC$ complex in DDM micelles. Species with an additional mass corresponding to copurified LPS are labelled with a star. For experimental details see the corresponding paragraph in the materials and methods section.

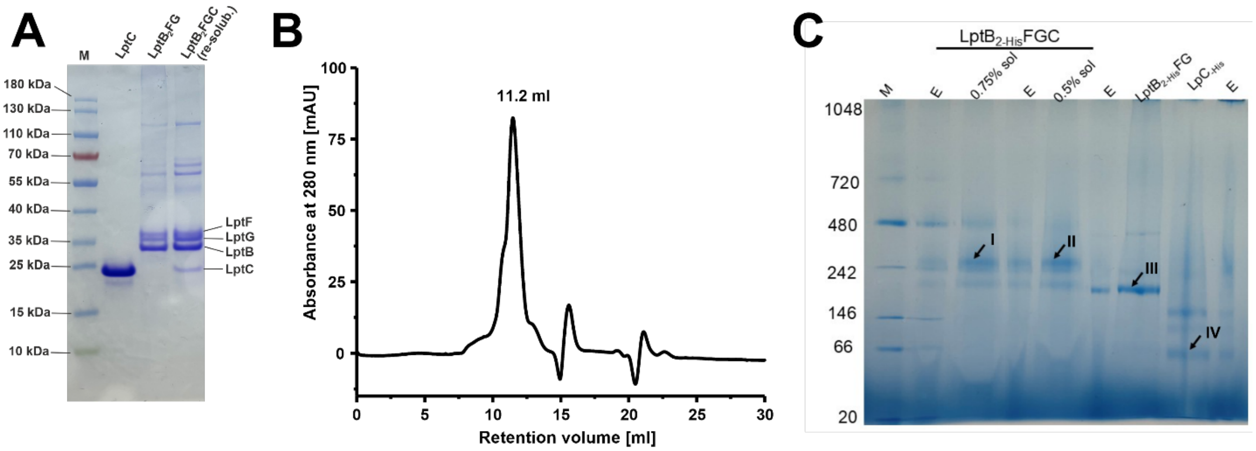


**Fig. S3:** Biochemical characterization of LptC, $LptB_2FG$ and $LptB_2FGC$. **(A)** Coomassie stained SDS gel of purified LptC, $LptB_2FG$ and re-solubilized $LptB_2FGC$ after *in vitro* complex formation. **(B)** Size exclusion chromatogram of purified LptC in detergent. The peak at 11.2 ml is composed of dimeric LptC (confirmed by native MS). **(C)** Blue-native PAGE gel of LptC, $LptB_2FG$ and $LptB_2FGC$. Labeled bands show I & II $LptB_2FGC$ complex, III $LptB_2FG$ complex and IV monomeric LptC.

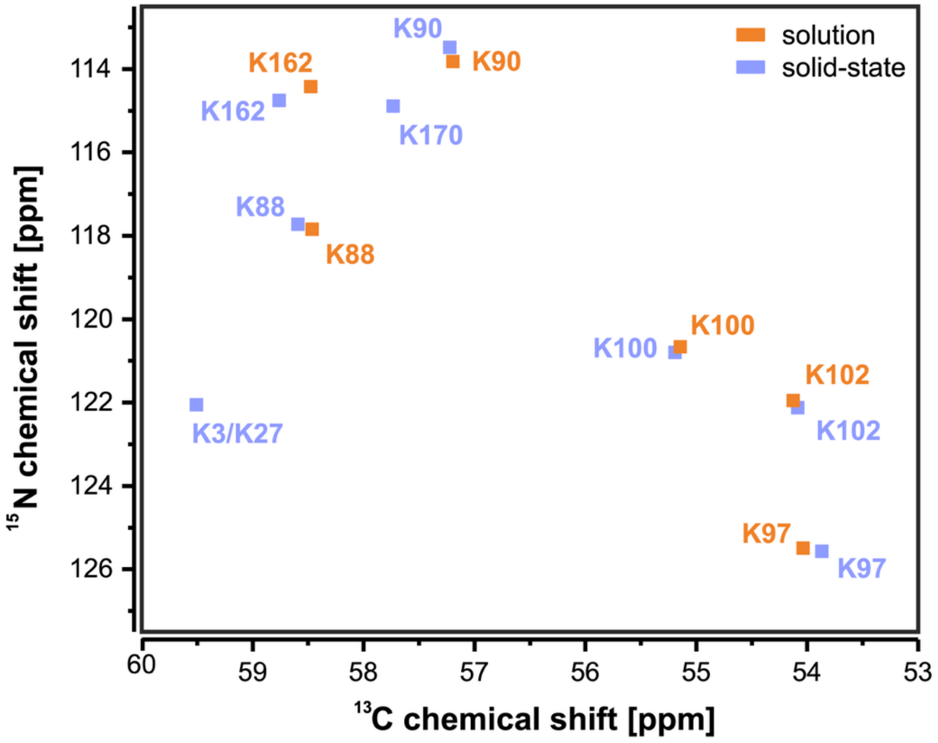


**Fig. S4:** Overlay of the 2D $^{13}$C, $^{15}$N DCP hNCA resonances of [$^{13}$C, $^{15}$N-K]-LptC in liposomes obtained here by ssNMR (blue) with the published resonance assignment (*1*) of the soluble ßJR domain of LptC (orange). K3/K27 and K170 were not assigned in solution (see materials and methods).

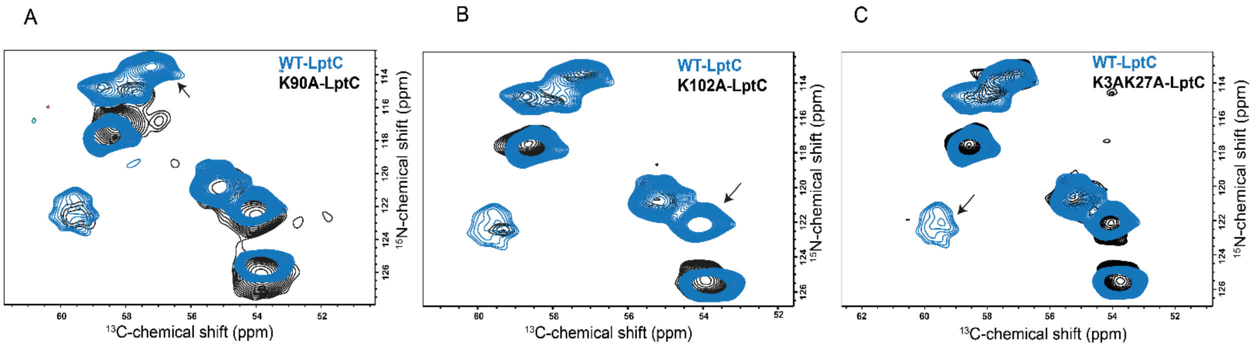


**Fig. S5:** hNCA ssNMR spectra of [$^{13}$C, $^{15}$N-K]-LptC and various LptC mutants for resonance assignment. **(A)** Comparison of LptC WT with LptC-K90A. The arrow points at the K90 resonance. **(B)** Comparison of LptC WT with LptC-K102A. The arrow highlights the disappearing resonance that can be assigned to K102. **(C)** Comparison of LptC WT with LptC-K3A-K27A. Both residues are located in the TMH and their overlapping signals disappear upon mutation (arrow).

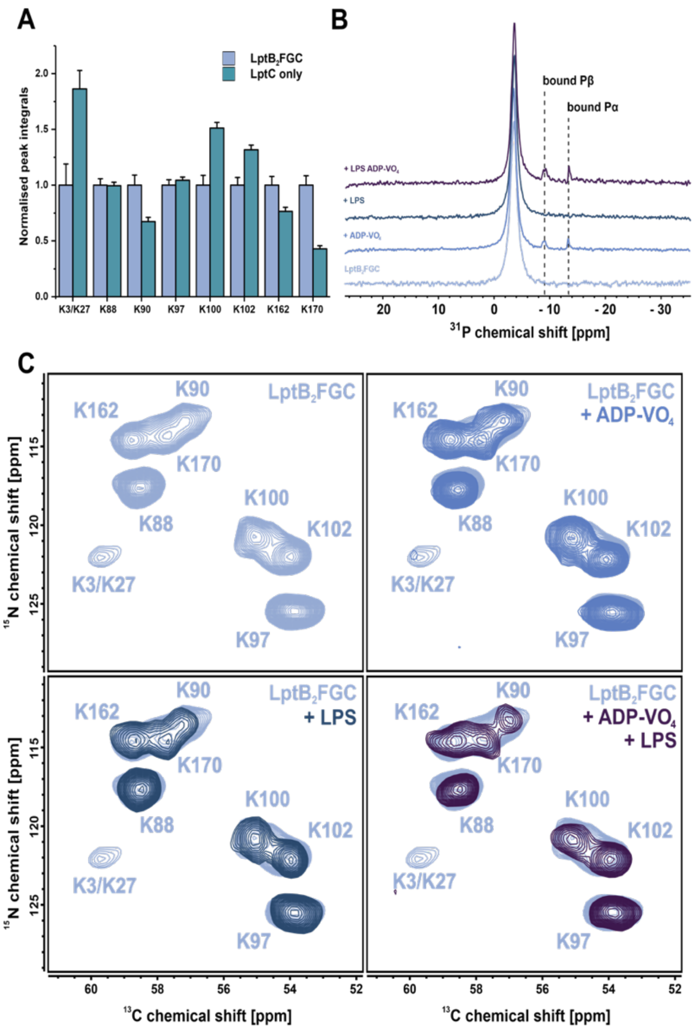


**Fig. S6:** Supporting NMR data. **(A)** Integrals corresponding to the volume of 2D cross peaks from lysines of only LptC alone derived from the spectra of Fig 3C. The integrals were normalized twice, first to the sum of all lysine integrals to correct the difference in sample amount and second to the integral of each peak of the apo $LptB_2FGC$ sample. The error bars represent the noise that contributes to the signal integral determined by the root mean square of noise-areas from the respective 2D spectra. **(B)** 1D $^{31}$P-CP MAS NMR spectra of the $LptB_2FGC$ used in the ambient temperature NMR experiments (Fig. 3) demonstrating successful trapping of the ADP.$VO_4$ states. Here, $^{1}$H-$^{31}$P cross polarization is used as a dynamic filter experiment visualizing only bound nucleotides. **(C)** Comparison of 2D $^{13}$C, $^{15}$N hNCA spectra of $LptB_2FG$ + [$^{13}$C, $^{15}$N-K]-C in liposomes in the apo state (top left) with the vanadate-trapped state (top right), the LPS-bound state (bottom left) and the vanadate-trapped state with LPS (bottom right). The TMH signal K3/K27 (59 ppm / 122 ppm) becomes weaker almost beyond detection in all states. Spectra are processed identically and show the same contour levels. All spectra were recorded with 14 kHz MAS spinning rate at an 850 MHz NMR spectrometer. For full experimental details see the corresponding paragraph in the materials and methods section.

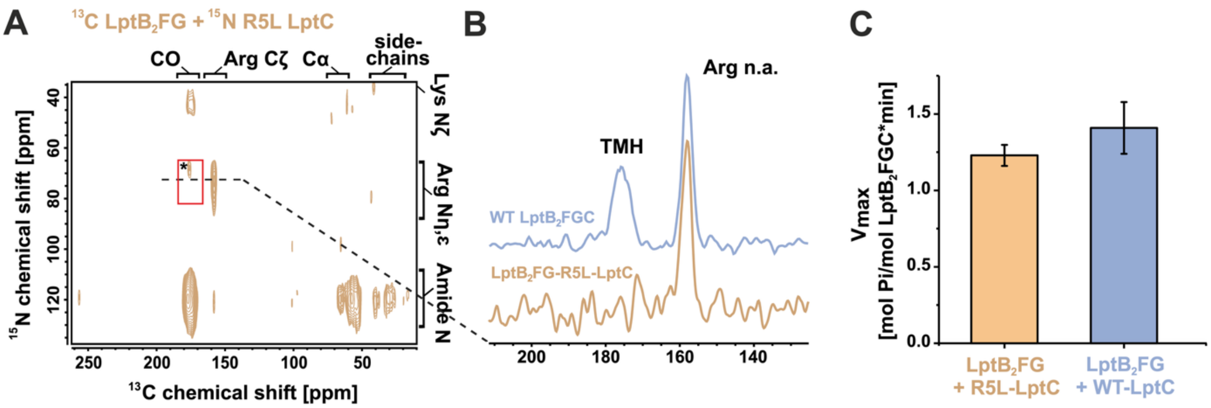


**Fig. S7:** Control experiments on $LptB_2FG$ in complex with LptC-R5L. **(A)** DNP-enhanced $^{13}C$, $^{15}N$-TEDOR spectra of the differentially labelled $LptB_2FG$ in complex with LptC-R5L in liposomes at 100 K in the apo state. The location of the specific LptC-LptF contact between Arg5 and Ser298 seen in the wild type (Fig. 4B) is indicated. **(B)** 1D $^{13}C$ slices through the expected cross peak region along $\omega_1(^{15}N)$ = 70 ppm in comparison with the wildtype complex. Upon introducing the LptC-R5L mutation, the signal intensity disappears allowing confident assignment to a specific LptC-LptF interaction. **(C)** ATPase activity of the $LptB_2FG$ complex with LptC-R5L is comparable to the wild type. Comparable $v_{max}$ values verify correct complex formation in both cases as shown on Fig. 2A. The molar $LptB_2FG$ to LptC ratio was 1:1.5.

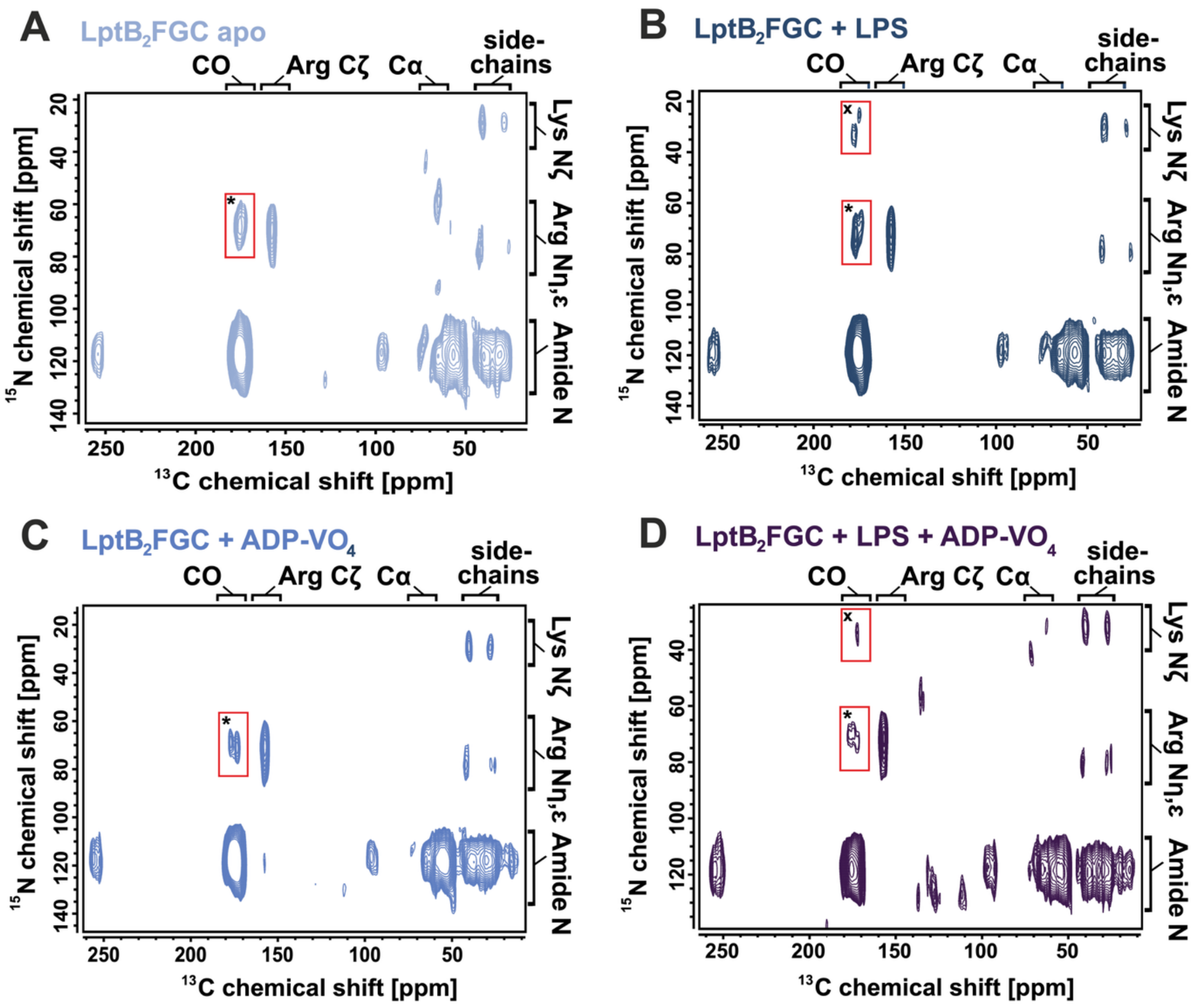


**Fig. S8:** DNP-enhanced $^{13}$C, $^{15}$N-TEDOR spectra of the differentially labelled LptB$_2$FGC complex in liposomes at 100 K in the **(A)** apo state, **(B)** LPS-bound state, **(C)** vanadate trapped state and **(D)** combined LPS-bound / vanadate trapped state. The intermolecular cross-peaks not arising from natural abundance contributions are highlighted by a red box. The assigned cross-contact of LptC R5-Nh,e with LptF S298-C' is labelled with (*). Interestingly, we observed another intermolecular contact (x) that is present only in the two LPS bound states. This intermolecular contact originates from a lysine sidechain nitrogen N$_\zeta$ of LptC and a backbone C' or Cγ/δ side chain atom of E, Q, D or N from LptB$_2$FG. The intensity of the cross peak is much lower compared to the TMH contact. The presence of LPS seems to introduce an additional contact site, which could originate from an interaction involving a Lys in LptC$_{TMH}$ or in LptC$_{\beta JR}$. For experimental details see the corresponding paragraph in the materials and methods section.

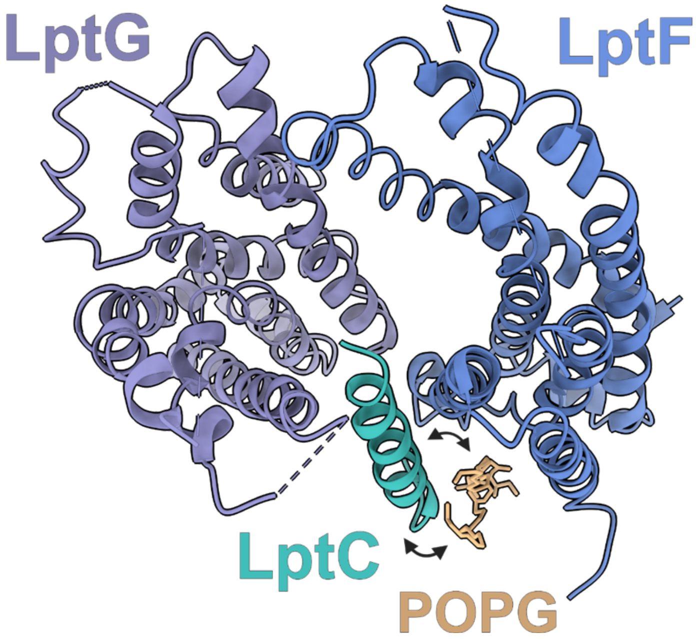


**Fig. S9:** Cryo-EM structure of $LptB_2FGC$ in nanodiscs (PDB:6MI7) shows a co-purified and bound POPG lipid density in proximity of the LptC transmembrane helix. For simplification only the TMDs of LptG, LptF and LptC are shown as top view. This binding site could serve as the proposed second low affinity binding site for the $LptC_{TMH}$, when dynamical exchange is triggered by LPS binding.

### Table S1. Primers for lysine mutants of LptC protein

| | | |
|---|---|---|
| **1** | K3A-LptC for<br>K3A-LptC rev | 5'-aacccaacgtctggctgcactcatcatatgtatatctccttcttaaagt-3'<br>5'-actttaagaaggagatatacatatgatgagtgcagccagacgttgggtt -3' |
| **2** | K27A-LptC for<br>K27A-LptC rev | 5'-ctgggcggtatcgtctgcttcggccatattaatgccg-3'<br>5'-cggcattaatatggccgaagcagacgataccgcccag-3' |
| **3** | K43A-LptC for<br>K43A-LptC rev | 5'-ccgtatgctcgcttgcataggtgggatcattgttgttgacga-3'<br>5'-tcgtcaacaacaatgatcccacctatgcaagcgagcatacgg-3' |
| **4** | K88A-LptC for<br>K88A-LptC rev | 5'-gtcgggattttatccgcatcaaacgtggtaagtaccggct-3'<br>5'-agccggtacttaccacgtttgatgcggataaaatcccgac-3' |
| **5** | K90A-LptC for<br>K90A-LptC rev | 5'-tttacggaccatgtcgggattgcatccttatcaaacgtggtaag-3'<br>5'-cttaccacgtttgataaggatgcaatcccgacatggtccgtaaa-3' |
| **6** | K97A-LptC for<br>K97A-LptC rev | 5'-agcttggctttatctgctgctacggaccatgtcgggatttt-3'<br>5'-aaaatcccgacatggtccgtagcagcagataaagccaagct-3' |
| **7** | K100A-LptC for<br>K100A-LptC rev | 5'-ggtcagcttggctgcatctgcttttacggaccatgtcgg-3'<br>5'-ccgacatggtccgtaaaagcagatgcagccaagctgacc-3' |
| **8** | K102A-LptC for<br>K102A-LptC rev | 5'- catccggtcattggtcagcgcggctttatctgcttttacg-3'<br>5'-cgtaaaagcagataaagccgcgctgaccaatgaccggatg -3' |
| **9** | K162A-LptC for<br>K162A-LptC rev | 5'-cgtaagttgccgcgcattgccagaccgctggagttaaa-3'<br>5'-tttaactccagcggtctggcaatgcgcggcaacttacg-3' |
| **10** | K170A-LptC for<br>K170A-LptC rev | 5'-aatcagctcggcgttcgcgctgcgtaagttgccg-3'<br>5'-cggcaacttacgcagcgcgaacgccgagctgatt-3' |
| **11** | K177A-LptC for<br>K177A-LptC rev | 5'- tcataggatgttctaaccgcttcaatcagctcggcgttcttg -3'<br>5'- caagaacgccgagctgattgaagcggttagaacatcctatga -3' |
| **12** | K187A-LptC for<br>K187A-LptC rev | 5'- gatgatgaggctgagtttgtgcgttttgaatttcataggatgttctaacc -3'<br>5'- ggttagaacatcctatgaaattcaaaacgcacaaactcagcctcatcatc-3' |
